# Development of a metabolomics-based index to monitor dietary effects on chronic inflammation: The Dietary Metabolomics Inflammation Index

**DOI:** 10.64898/2026.07.06.736618

**Authors:** Jiada Zhan, Chin-An Yang, Mary Nellis, Youran Tan, Matthew Ryan Smith, Jessica Alvarez, Donghai Liang, Anne Dunlop, Greg Martin, Young-Mi Go, Dean P. Jones

## Abstract

**Background:** The Dietary Inflammatory Index (DII) is widely used to assess the inflammatory potential of diet, but it relies on self-reported dietary assessment and does not directly capture individual differences in metabolism as an intermediate connection to inflammation. High-resolution metabolomics provides objective measurements that complement dietary assessment to support precision nutrition to control inflammation.

**Objective:** We developed, tested, and applied a Dietary Metabolite Inflammatory Index (DMII) to assess diet-related chronic inflammation using metabolites measured by liquid chromatography–high-resolution mass spectrometry.

**Methods:** DII was calculated using *dietaryindex* R package with Block Food Frequency Questionnaire (FFQ) data. To develop the DMII, chronic inflammation-related dietary metabolites corresponding to the DII food parameters were found through a literature review. Dietary metabolites were identified and quantified by authentic standards by our established laboratory procedures. DMII uses the same inflammatory effect scores as the DII. Three DMII versions were developed: concentration-based, median-based, and quintile-based DMII. Mean and standard deviation of 29 dietary metabolites were calculated by using 3025 human plasma samples from 3 studies. DMII was tested in the Center for Health Discovery and Well-Being cohort (CHDWB) and the Atlanta African American Maternal and Child cohort (ATLAA) using chronic inflammation biomarkers, including high-sensitivity C-reactive protein (hs-CRP), CRP, and IL-6. The median-based DMII was further applied to four Alzheimer’s disease metabolomics datasets as a proof-of-concept application.

**Results:** In the CHDWB study, concentration-based DMII had a weak positive correlation with Block FFQ-derived DII and strongly correlated with median-based and quintile-based DMII. In the same study, all three DMII versions had significant positive correlations with hs-CRP and IL-6. In the ATLAA study, only concentration-based DMII was positively associated with CRP and IL-6. Higher median-based DMII was associated with higher odds of Alzheimer’s disease.

**Conclusions:** DMII provides a metabolomics-based framework for assessing diet-related chronic inflammation using metabolomics data. This metabolomics approach may complement self-reported dietary assessment to use diet and nutrition to help protect against chronic disease linked to inflammation.

## Introduction

Dietary patterns capture the combined effects of multiple foods and nutrients. They are therefore widely used to study associations between diet and chronic disease (1,2). Several dietary pattern indexes have been developed, including the Healthy Eating Index 2020 (3), Alternative Healthy Eating Index (4), Dietary Approach to Stop Hypertension Index (DASH) (5), Alternate Mediterranean Diet Score (aMED) (6), and Dietary Inflammatory Index (DII) (7). These indexes have improved our understanding of diet–disease relationships and have helped guide strategies for disease prevention and treatment.

Among these indexes, the DII has been widely used to assess the inflammatory potential of diet. A higher DII score indicates consumption of a more pro-inflammatory diet, whereas a lower DII score is reflective of a more anti-inflammatory diet. This framework is biologically relevant because chronic inflammation contributes to many chronic diseases, including cardiovascular diseases (8), type 2 diabetes (9), cancers (10), obesity (11), and Alzheimer’s disease (12). Consistently, studies using the DII have linked pro-inflammatory dietary patterns to these diseases (13–16).

Dietary pattern index analysis relies on standardized dietary data. These data are usually collected using food frequency questionnaires, 24-hour dietary recalls, or food records (2). Although these methods have been extensively validated, they do not fully capture individual differences in the absorption, distribution, metabolism, and elimination of foods and nutrients. They are also affected by misreporting, which is a well-known limitation of self-reported dietary assessment (17,18). These limitations are especially important for precision nutrition, which aims to understand how individuals differ in their responses to foods and nutrients (19).

Metabolomics offers a complementary approach to dietary assessment. It provides a systems-level link among the genome, environmental exposures, and health-related phenotypes in physiology and disease (20,21). Liquid chromatography-high resolution mass spectrometry (LC-HRMS), in particular, enables high-throughput untargeted metabolomic profiling of tens of thousands of metabolic features (22,23). This approach can identify measurements of dietary exposure in human biospecimens such as blood, urine, and cerebrospinal fluid (24–27). Therefore, dietary metabolites can be useful for precision nutrition because they reflect the absorption, metabolism, and disposition of foods and nutrients (24). Dietary metabolite can serve as objective measurements of food intake (24,25). These measurements may better capture the physiological effects of diets compared to self-reported dietary intake alone (26). Dietary metabolites can therefore complement traditional dietary assessments. In some settings, they may also provide an alternative when dietary data are unavailable or when more precise nutritional measurements are needed (26). This concept was described as nutritional metabolomics that uses metabolite profiling in a global manner to study diet and health (24), and it was demonstrated in previous studies (26–29).

Here, we integrate the concepts of the DII and LC-HRMS untargeted metabolomics to develop a Dietary Metabolite Inflammatory Index (DMII). The DMII uses literature-supported dietary metabolites linked to DII food and nutrient parameters to assess diet-related chronic inflammation with a standardized scoring system. Either metabolite concentrations or raw metabolite intensities can be used to calculate the DMII. This approach provides a convenient and objective measurement of diet-related chronic inflammation for individuals and study populations. As the metabolomics data become increasingly available, the DMII can expand the study of diet-related inflammation in cohorts that lack traditional dietary assessment and support studies of biological mechanisms research related to inflammation.

## Methods

### Development of the DMII

To develop the DMII, we used DII as the conceptual framework (7). The DII is an established dietary index that assesses individual’s diet-related chronic inflammation using 45 food and nutrient parameters with a standardized scoring system (7). It first developed a global nutrition database for the literature-supported 45 inflammation-related food parameters. This generated the average and standard deviation of the food intake of the DII food parameters. Then, through literature review, each DII food parameter has its own weight, or called “overall food parameter-specific inflammatory effect score” by summarizing the type and number of literatures that link inflammation and the food parameter. Later, in each study, dietary data from individuals is used to calculate Z-score and centered percentiles for each of the food parameters using the global average and standard deviation. The centered percentile value for each food parameter is then multiplied by the corresponding weight to obtain the DII score specific to the food parameter. Lastly, all food parameter-specific DII scores are sum up to obtain the total DII score for an individual.

We first reviewed the HMDB database to identify known dietary metabolites that correspond to the DII food parameters (30). This enabled us to identity many well-documented metabolites related to the DII food parameters, such as vitamin B6 intake and pyridoxal, caffeine intake and caffeine, monounsaturated fatty acid intake and palmitoleic acid, oleic acid, and gondoic acid (**Table 1 and Supplementary Table 1**). We then reviewed the literature to see if the other DII food parameters are linked with metabolites (**Table 1 and Supplementary Table 1**). Finally, only metabolites that were identified and quantified in blood by our established laboratory procedures were considered for the DMII (31).

**Table 1.** Concentration-based Dietary Metabolite Inflammatory Index.

| Food parameter | Metabolite parameter | Overall inflammatory effect score | Mean metabolite concentration (μmol/L) | Standard deviation of metabolite concentration |
| --- | --- | --- | --- | --- |
| Vitamin B6 | Pyridoxal (B6) | -0.365 | 0.03 | 0.03 |
| Caffeine | Caffeine | -0.110 | 4.07 | 8.73 |
| Carbohydrate | Glucose | 0.097 | 6678.38 | 2865.06 |
| Cholesterol | Cholesterol | 0.110 | 1234.34 | 580.70 |
| Fiber | Indole propionate | -0.663 | 10.04 | 24.79 |
| Green/black tea | Hippuric acid | -0.536 | 6.05 | 9.11 |
| MUFA | Palmitoleic acid | -0.009 | 13.44 | 14.55 |
|  | FA 18:1 (Oleic acid) |  | 176.40 | 200.47 |
|  | FA 20:1 (Gondoic acid) |  | 34.52 | 51.96 |
| Niacin | Nicotinamide (B3) | -0.246 | 1.12 | 1.10 |
| n-3 fatty acid | FA 20:5 (Eicosapentaenoic acid) | -0.436 | 51.44 | 66.95 |
|  | DHA (Docosahexaenoic acid) |  | 47.25 | 44.00 |
| n-6 fatty acid | FA 20:4 (Arachidonic acid) | -0.159 | 998.75 | 1001.89 |
|  | FA 18:2 (Linoleic acid) |  | 5637.75 | 5643.27 |
|  | FA 20:3 (Homolinoleic acid) |  | 145.31 | 133.46 |
|  | FA 20:2 (Eicosadienoic acid) |  | 21.21 | 19.94 |
|  | FA 22:4 (Docosatetraenoic acid) |  | 23.19 | 15.69 |
| Protein | Delta-valerobetaine | 0.021 | 1.62 | 2.38 |
| PUFA | Gamma-Linolenic acid and Alpha-Linolenic acid | -0.337 | 82.12 | 94.31 |
|  | Docosapentaenoic acid (22n-3) and Docosapentaenoic acid (22n-6) |  | 52.27 | 44.93 |
| Saturated fat | FA 14:0 (Myristate) | 0.373 | 7.85 | 7.13 |
|  | FA 16:0 (Palmitate) |  | 88.33 | 56.77 |
|  | FA 17:0 (Heptadecanoic) |  | 2.26 | 2.20 |
|  | FA 18:0 (Stearic acid) |  | 61.51 | 31.06 |
|  | FA 20:0 (Arachidic acid) |  | 22.50 | 31.47 |
| Vitamin A | Retinol | -0.401 | 7.02 | 3.59 |
| Vitamin C | Ascorbate | -0.424 | 834.96 | 1522.55 |

MUFA Monounsaturated fatty acids, PUFA Polyunsaturated fatty acids. Metabolites included in the Dietary Metabolite Inflammatory Index are identified by linking foods and nutrients in the Dietary Inflammatory Index with the associated metabolites through a literature review. Overall inflammatory effect scores are from the Dietary Inflammatory Index. Metabolite concentration mean and standard deviation values are from the quantification results of 3025 plasma samples across 3 human studies analyzed by the same liquid chromatography mass spectrometry setting.

In general, DMII follows a similar calculation method and uses the same weights adopted in the DII food parameters. For different scenarios, we developed three versions of DMII: concentration-based DMII, median-based DMII, and quintile-based DMII (**Figure 1**). The concentration-based DMII requires metabolite identification and quantification. In contrast, the median-based and quintile-based DMII require metabolite identification but not absolute quantification. All three versions used the same metabolite components and overall inflammatory effect scores (weights).

**Figure 1.**
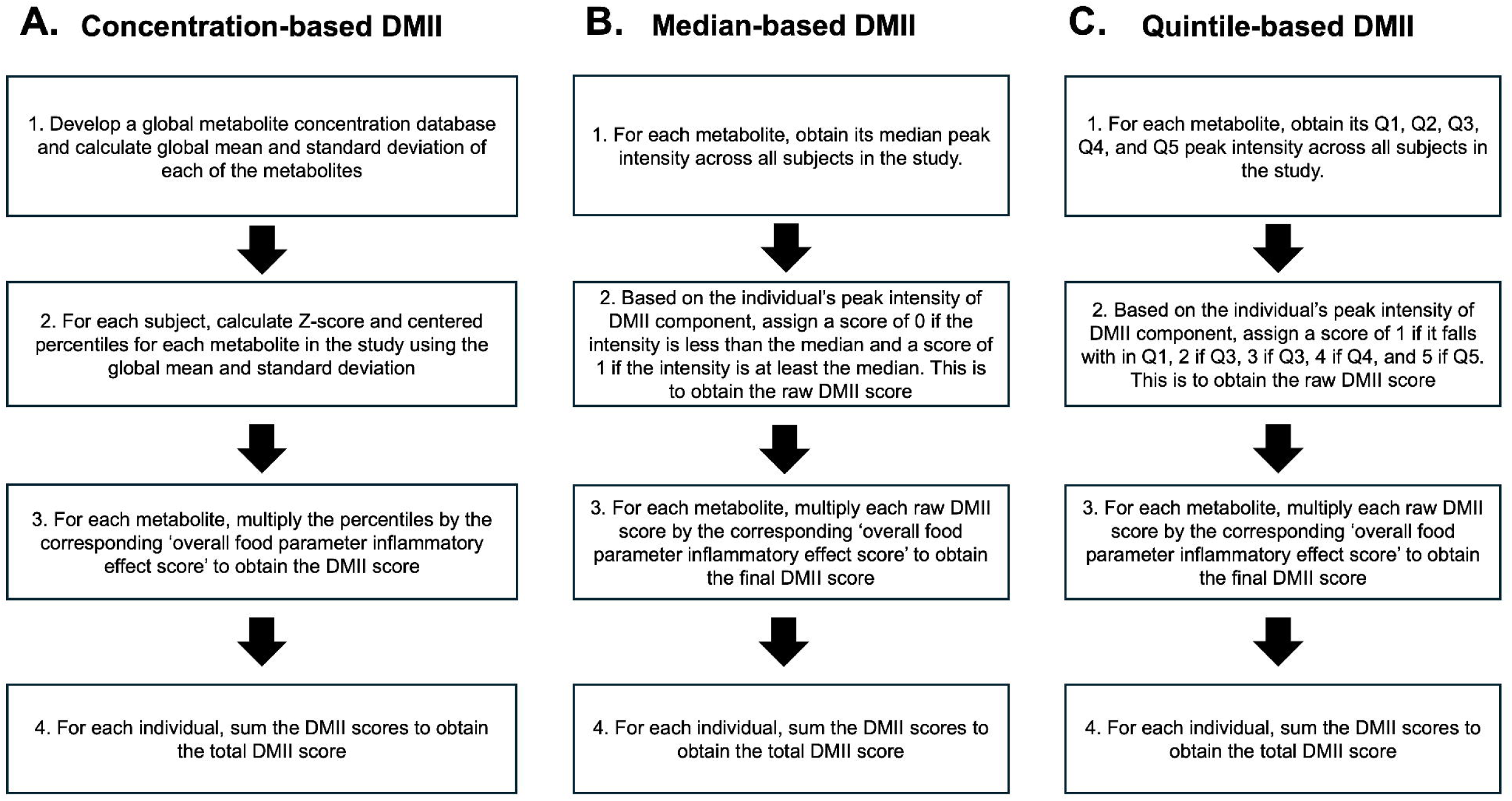
Flowchart of Dietary Metabolite Inflammatory Index calculation. For all versions of the Dietary Metabolite Inflammatory Index (DMII), metabolite identification is necessary. For concentration-based DMII, metabolite quantification is also needed. For median- and quintile-based DMII, if multiple metabolites are identified for the same DMII component, their intensities are summed to obtain the total intensities, which are then used to calculate the median and quintiles.

Concentration-based DMII is the most similar form of DMII compared to DII (**Figure 1A**). It first develops a global metabolite concentration database for the literature-supported 29 metabolites related to DII food parameters. The database was built from the metabolite concentrations in 3 in-house reference human plasma metabolomics studies that included 3025 samples. Some of the samples were from the same subjects with repeated measures. This generates the average and standard deviation of metabolite concentration of the DMII metabolite parameters in the global metabolite concentration database. Meanwhile, each DMII metabolite parameter uses the same weights as the relevant DII food parameters. Then, in each study, metabolite concentrations from individuals is used to calculate Z-score and centered percentiles for each of the DMII metabolite parameters using the global average and standard deviation. The centered percentile value for each metabolite parameter is then multiplied by the corresponding weight to obtain the DMII score specific to the metabolite parameter. Lastly, all metabolite parameter-specific DMII scores are sum up to obtain the total DMII score for an individual.

The median-based DMII is adapted from the median-based scoring design of the aMED index and uses the same weights used in the DII (6,7) (**Figure 1B**). For each DMII component, participants with metabolite intensity above the study-specific median received a raw score of 1. Participants with metabolite intensity at or below the median received a raw score of 0. Each DMII raw component score was then multiplied by the corresponding weights. The weighted component scores were summed to calculate the total median-based DMII score.

The quintile-based DMII is adapted from the quintile-based scoring design of the DASH index and uses the same weights used in the DII (5,7) (**Figure 1C**). For each DMII component, participants were ranked by metabolite intensity and classified into study-specific quintiles. Participants in quintile 1 received a raw score of 1, and participants in quintile 5 received a raw score of 5. Each DMII raw component score was multiplied by the corresponding weights. The weighted component scores were summed to calculate the total quintile-based DMII score.

### Study population

Two metabolomics studies were used to test the DMII. The first validation study included 212 plasma samples with both metabolomics data and inflammatory biomarkers from adults in the Center for Health Discovery and Well-Being (CHDWB) cohort (32). Participants were classified as metabolically healthy nonobese (MHNO), metabolically healthy obese (MHO), or metabolically unhealthy obese (MUO). The second validation study included 598 serum samples with both metabolomics data and inflammatory biomarkers from two clinic visits of pregnant African American women in the prospective Atlanta African American Maternal and Child cohort (ATLAA) cohort (33,34).

Three external metabolomics studies were used to demonstrate the application of the DMII. These studies included Alzheimer’s disease (AD) datasets from three published studies: the EFIGA and WHICAP plasma datasets from the AD Knowledge Portal (https://www.synapse.org/Synapse:syn70083418/wiki/635413)(35), and paired plasma and cerebrospinal fluid datasets from study ST000046 in the Metabolomics Workbench (36). These datasets include subjects with Alzheimer’s disease and subjects as healthy controls. The EFIGA dataset included 150 patients with AD and 567 healthy controls. The WHICAP dataset included 100 patients with AD and 251 healthy controls. The ST000046 study included 15 patients with AD and 15 healthy controls.

### Health and dietary assessment

In the CHDWB study, chronic inflammation was assessed using high-sensitivity C-reactive protein (hs-CRP), IL-6, and TNF-α. Age, race, and sex were self-reported. Detailed health assessment procedures have been published elsewhere (32).

In the ATLAA study (33), maternal serum samples collected at the 8–14 weeks visit and 24-30 weeks visit were analyzed for 598 participants by the Emory Multiplexed Immunoassay Core to quantify IL-6 and CRP. IL-6 was measured using a Meso Scale Discovery assay platform (Meso Scale Diagnostics, Rockville, MD), which uses electrochemiluminescence detection to provide high sensitivity and a broad dynamic range, following the manufacturer’s protocol (37,38). CRP was measured with an enzyme-linked immunosorbent assay kit from R&D Systems (cat. no. SCRP00), also according to the manufacturer’s instructions. CRP concentrations were determined from four-parameter calibration curves generated in BioTek Gen5 software.

Dietary intake was assessed using the Block Food Frequency Questionnaire (FFQ) in both the CHDWB and ATLAA studies (39–41). DII was calculated using the *dietaryindex* R package, a validated tool for dietary index calculation (2).

For the DMII application analyses, AD diagnosis was defined according to the criteria used in each original study. Metadata for these studies were downloaded from the AD Knowledge Portal and the Metabolomics Workbench.

### Metabolomics data

For the 3 reference studies, the CHDWB study, and the ATLAA study (42,43), human plasma or serum samples were analyzed by LC-HRMS using established protocols (31). Samples were snap-frozen and maintained at -80 °C before the analysis. The LC system was a Thermo Scientific Dionex Ultimate 3000, coupled either to a Thermo Scientific HF-Q Exactive Fourier transform high-resolution mass spectrometer or to a Thermo Fisher Orbitrap Fusion Tribrid instrument. The chromatographic platform used a dual-pump configuration, which enabled parallel analyte separation and column washing.

Sample extracts were analyzed using two complementary LC-HRMS methods: Hydrophilic Interaction Chromatography in positive ion mode (HILIC+) and Reverse Phase in negative ion mode (C18-). The HILIC+ method used a Waters XBridge BEH Amide XP HILIC column (2.1 × 50 mm, 2.6 μm particle size) with mobile phase A consisting of LC-HRMS-grade water, mobile phase B consisting of LC-HRMS-grade acetonitrile, and mobile phase C consisting of 2% formic acid. The gradient was held at 22.5% A, 75% B, and 2.5% C for 1.5 min, then changed linearly to 75% A, 22.5% B, and 2.5% C by 4 min, followed by a 1-min hold.

The C18- method used a Higgins Targa C18 column (2.1 × 50 mm, 3 μm particle size) with mobile phase A consisting of water, mobile phase B consisting of acetonitrile, and mobile phase C consisting of 10 mM ammonium acetate. The gradient was held at 60% A, 35% B, and 5% C for 1 min, then changed linearly to 0% A, 95% B, and 5% C over the next 3 min, followed by a 2-min hold. For both methods, the flow rate was 0.35 mL/min during the first minute and was then increased to 0.4 mL/min for the remaining 4 min.

Each sample was analyzed by LC-HRMS as triplicates. Metabolomics features were extracted by apLCMS (44) and xMSanalyzer (45). Feature intensities were median-summarized and batch-corrected using the Combat algorithm (45).

For the concentration-based DMII, metabolites were identified as Schymanski Level 1 identifications by matching mass-to-charge ratio (± 10 ppm), retention time (± 30 s), and ion dissociation spectra (MS/MS) of study features to authentic chemical standards (**Supplementary Table 1**) (46). Metabolites were then quantified using a reference standardization approach (31,47). The metabolite concentration unit is μmol/L (μM).

For the median-based and quintile-based DMII, metabolites were first identified using the same Schymanski Level 1 identification method and then identified using the MSMICA algorithm. When metabolomics features or metabolite identities were in conflict between the reference library results and MSMICA results, the reference library results were prioritized. MSMICA is a recently developed metabolite identification algorithm that does not require chemical standards or MS/MS spectra (48). Instead, it integrates multiple sources of LC-HRMS and biological evidence to support metabolite identification and helps identify metabolites when their retention times are outside of the retention time matching window (30 seconds).

### Statistical analysis

Demographic characteristics were summarized using the TernTables R package (49). Descriptive statistics and univariate comparisons were used to compare participant characteristics across study groups.

Spearman correlations were used to evaluate agreement between the DII and each version of the DMII in the CHDWB study. Spearman correlations were also used to test the DMII by testing associations of the DII and each DMII version with inflammatory biomarkers in the CHDWB and ATLAA studies. Analyses were performed using complete observations for each inflammatory biomarker. Statistical significance was defined as p < 0.05.

As a proof-of-concept application, logistic regression was used to test whether the median-based DMII differed between patients with Alzheimer’s disease (AD) and healthy controls. This is because metabolites were not quantified, and there are only 15 AD patients and 15 healthy controls in the ST000046 studies, so using median-based DMII may improve the reliability of the statistics. Age and sex were adjusted in the EFIGA and WHICAP studies. The ST000046 study was not adjusted for age or sex because these variables were not available. Odds ratios (ORs) were scaled to each study-specific standard deviation of the median-based DMII. This was done by multiplying the logistic regression coefficient and its standard error by the standard deviation of the median-based DMII in each study. The rescaled coefficient was then exponentiated. Therefore, each OR represents the association with Alzheimer’s disease per one study-specific standard deviation higher median-based DMII.

Study-specific log ORs were pooled using inverse-variance weighting. Weights were calculated as the inverse of the squared standard error. The pooled log OR was then transformed back to the odds ratio scale to obtain the summary OR. The 95% confidence interval was calculated from the pooled standard error.

## Results

### DMII calculation

Using the metabolite reference standardization workflow (31,47), we quantified 29 DMII-related metabolites in 3025 human plasma samples from 3 studies (**Supplementary Table 1)**. These metabolites were measured using the HILIC+ method (**Figure 2**) and the C18- method (**Figure 3**). Most metabolite concentration distributions were skewed, supporting the use of Spearman correlation for later analyses.

**Figure 2.**
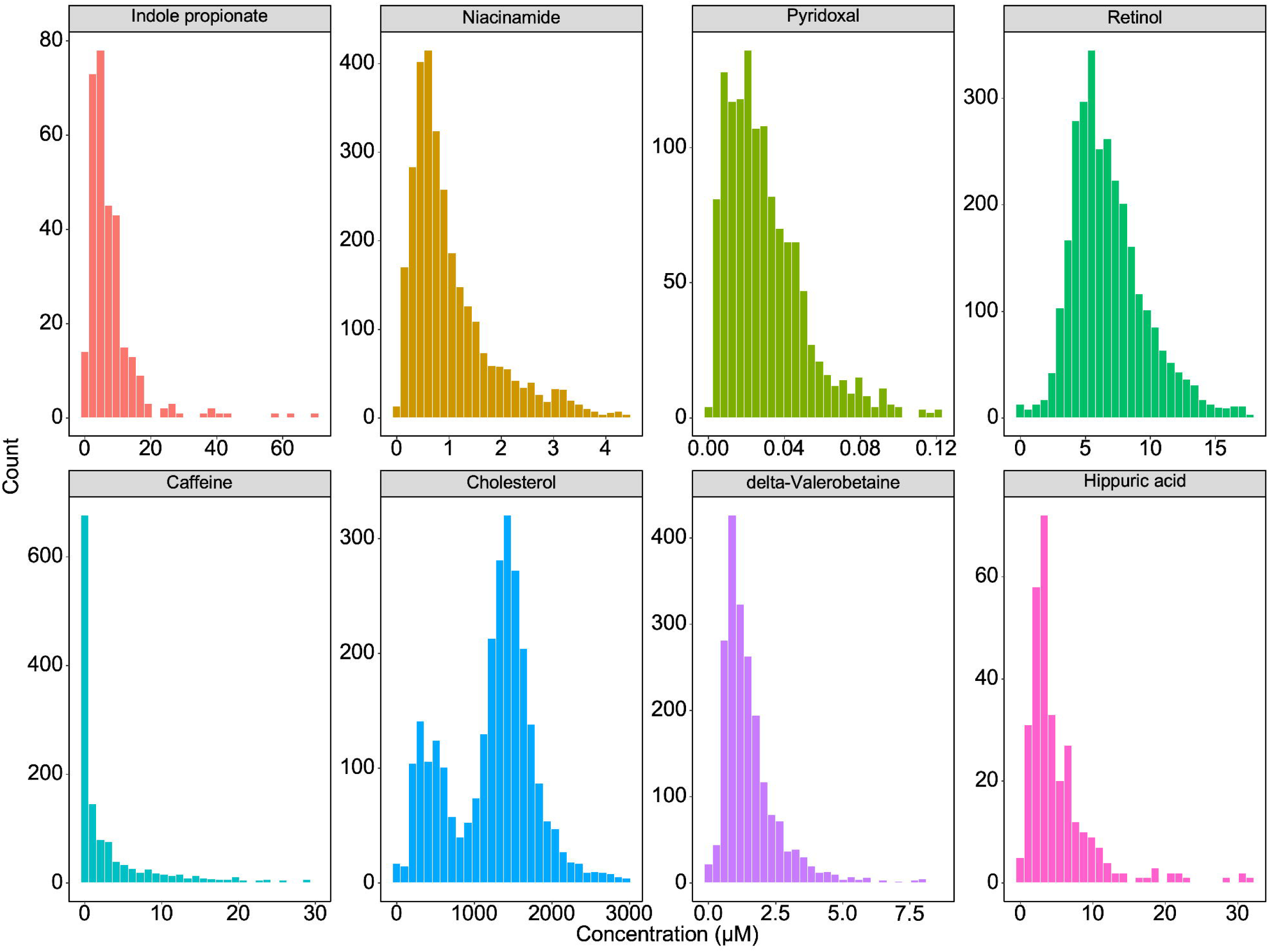
Histogram of the global HILIC positive metabolite concentrations. Metabolites were quantified using the reference standardization approach in the HILIC positive metabolomics data among the 3025 plasma samples. For the histogram of the metabolite concentrations, the metabolite concentrations of 0 μM and extreme outliers (greater than 3 standard deviations) were excluded to improve visualization.

**Figure 3.**
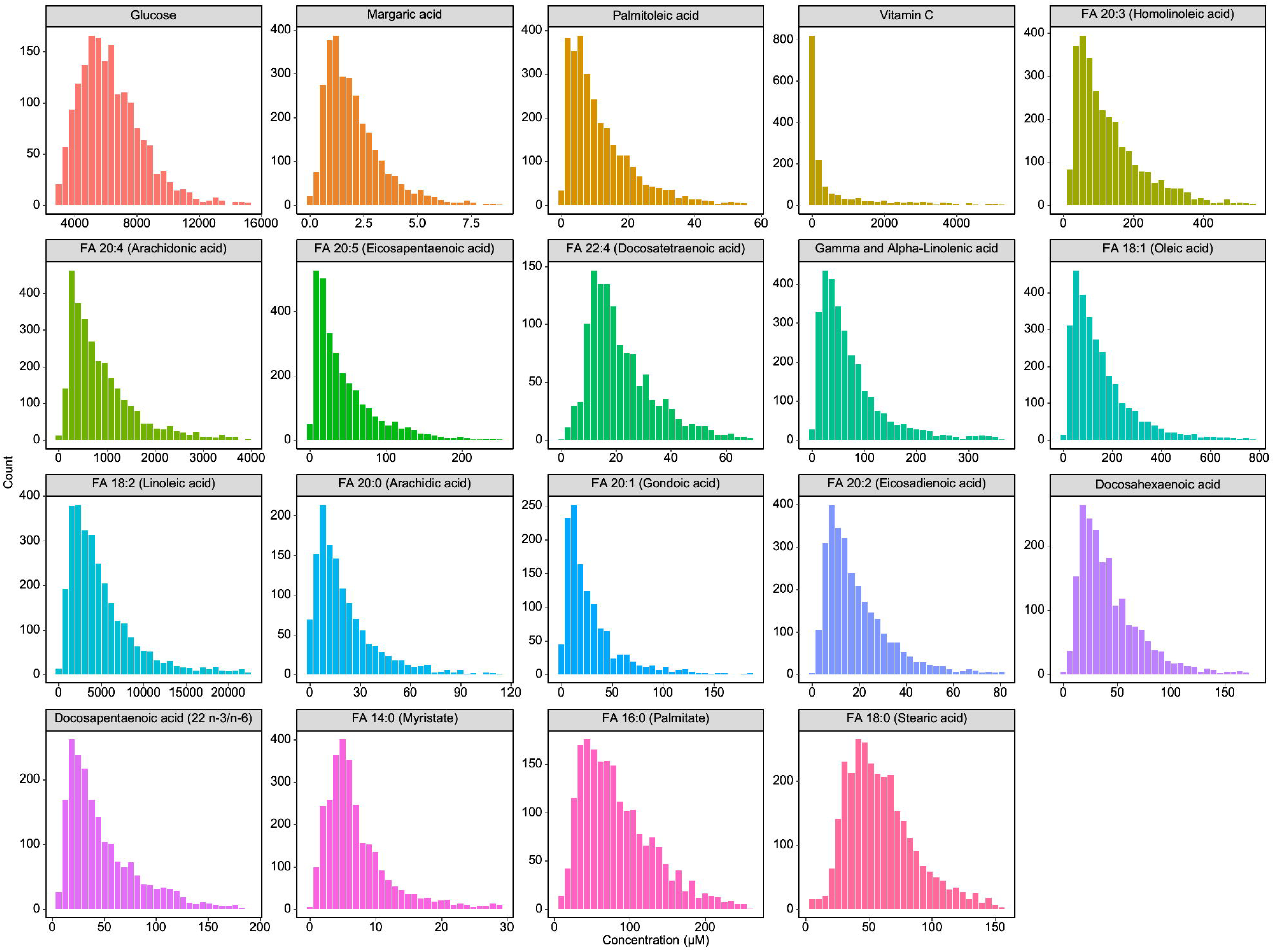
Histogram of the global C18 negative metabolite concentrations. Metabolites were quantified using the reference standardization approach in the C18 negative metabolomics data among 3025 plasma samples. For the histogram of the metabolite concentrations, the metabolite concentrations of 0 μM and extreme outliers (greater than 3 standard deviations) were excluded to improve visualization.

The global means and standard deviations of these metabolite concentrations were used as reference values for the concentration-based DMII (**Table 1 and Supplementary Table 1**). As described previously, the median-based DMII (**Table 2**) and quintile-based DMII (**Table 3**) were developed for settings in which metabolites are identified but not quantified.

**Table 2.** Median-based Dietary Metabolite Inflammatory Index.

| Food parameter | Metabolite parameter | Inflammatory effect | Raw Score of 0 | Raw Score of 1 | Overall inflammatory effect score | Final inflammatory score |
| --- | --- | --- | --- | --- | --- | --- |
| Vitamin B6 | Pyridoxal (B6) | anti-inflammatory | < median | >= study median | -0.365 | Individual raw score * individual inflammatory effect score |
| Caffeine | Caffeine | anti-inflammatory | < median | >= study median | -0.110 |  |
| Carbohydrate | Glucose | pro-inflammatory | < median | >= study median | 0.097 |  |
| Cholesterol | Cholesterol | pro-inflammatory | < median | >= study median | 0.110 |  |
| Fiber | Indole propionate | anti-inflammatory | < median | >= study median | -0.663 |  |
| Green/black tea | Hippuric acid | anti-inflammatory | < median | >= study median | -0.536 |  |
| MUFA | Palmitoleic acid<br>FA 18:1 (Oleic acid)<br>FA 20:1 (Gondoic acid) | anti-inflammatory | < median | >= study median | -0.009 |  |
| Niacin | Nicotinamide (B3) | anti-inflammatory | < median | >= study median | -0.246 |  |
| n-3 fatty acid | FA 20:5 (Eicosapentaenoic acid)<br>DHA (Docosahexaenoic acid) | anti-inflammatory | < median | >= study median | -0.436 |  |
| n-6 fatty acid | FA 18:2 (Linoleic acid)<br>FA 20:2 (Eicosadienoic acid)<br>FA 20:3 (Homolinoleic acid)<br>FA 20:4 (Arachidonic acid)<br>FA 22:4 (Docosatetraenoic acid) | anti-inflammatory | < median | >= study median | -0.159 |  |
| Protein | Delta-valerobetaine | pro-inflammatory | < median | >= study median | 0.021 |  |
| PUFA | Gamma-Linolenic acid<br>Alpha-Linolenic acid<br>Docosapentaenoic acid (22n-3)<br>Docosapentaenoic acid (22n-6) | anti-inflammatory | < median | >= study median | -0.337 |  |
| Saturated fat | FA 14:0 (Myristate)<br>FA 16:0 (Palmitate)<br>FA 17:0 (Heptadecanoic)<br>FA 18:0 (Stearic acid)<br>FA 20:0 (Arachidic acid) | pro-inflammatory | < median | >= study median | 0.373 |  |

| Table 2. Median-based Dietary Metabolite Inflammatory Index |  |  |  |  |  |  |
| --- | --- | --- | --- | --- | --- | --- |
| Food parameter | Metabolite parameter | Inflammatory effect | Raw Score of 0 | Raw Score of 1 | Overall inflammatory effect score | Final inflammatory score |
| Vitamin A | Retinol | anti-inflammatory | < median | >= study median | -0.401 |  |
| Vitamin C | Ascorbate | anti-inflammatory | < median | >= study median | -0.424 |  |

Median-based Dietary Metabolite Inflammatory Index uses the median intensity of identified metabolites for estimating dietary inflammation. When multiple metabolites are available for one food parameter, their LC-HRMS intensities are summed up to obtain the total intensity. The total intensity of each food parameter is then used to calculate the DMII raw score. For each DMII food parameter, its associated metabolite intensity from all participants is used to calculate the median intensity for the parameter. When one participant's metabolite intensity of a specific food parameter is lower than the median intensity, then its raw score is 0. The final inflammatory score of each DMII component is derived from multiplying the raw score by the overall inflammatory effect score. In the end, the final inflammatory scores of all DMII components are summed up to obtain the total final inflammatory score.

**Table 3.**
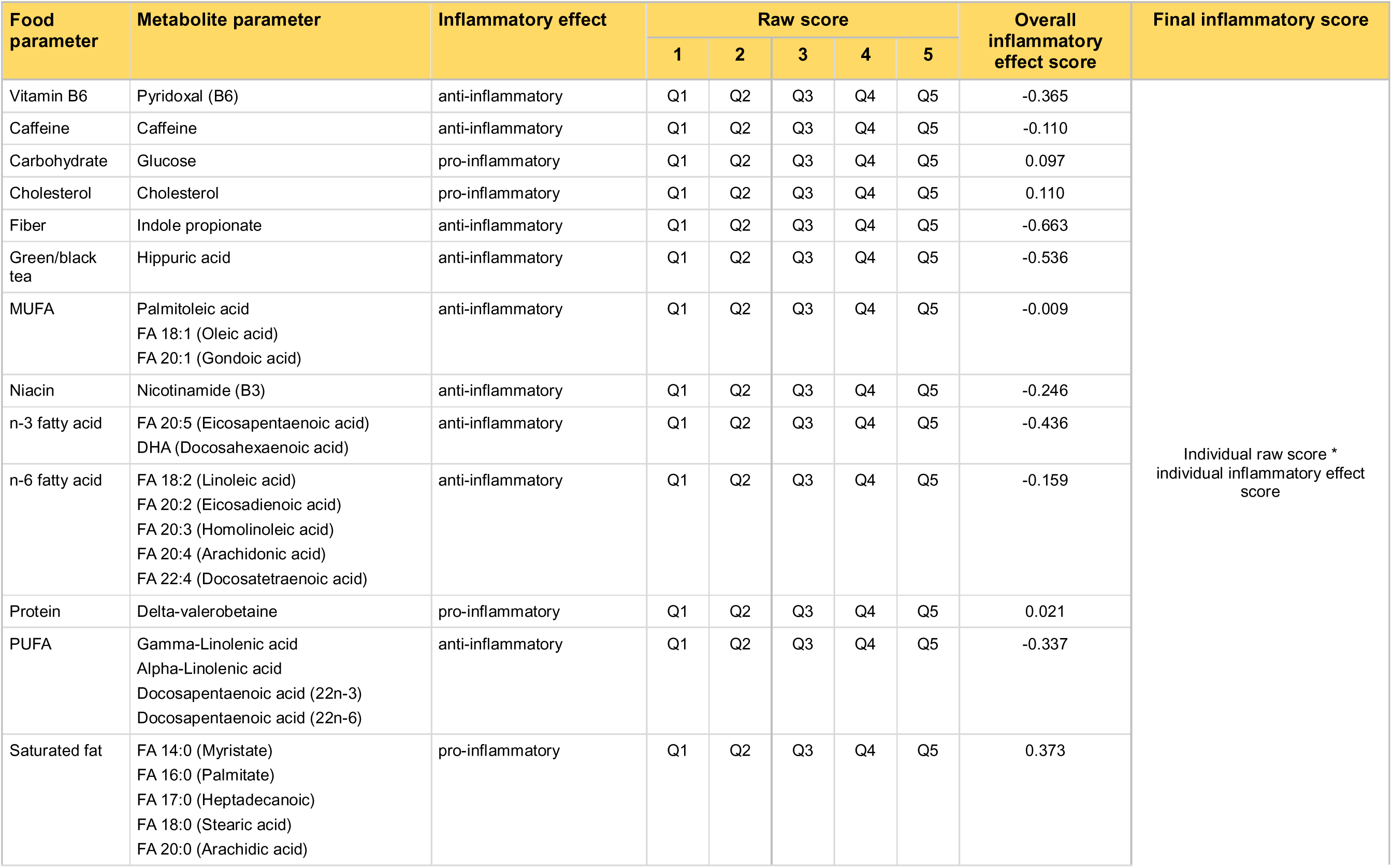
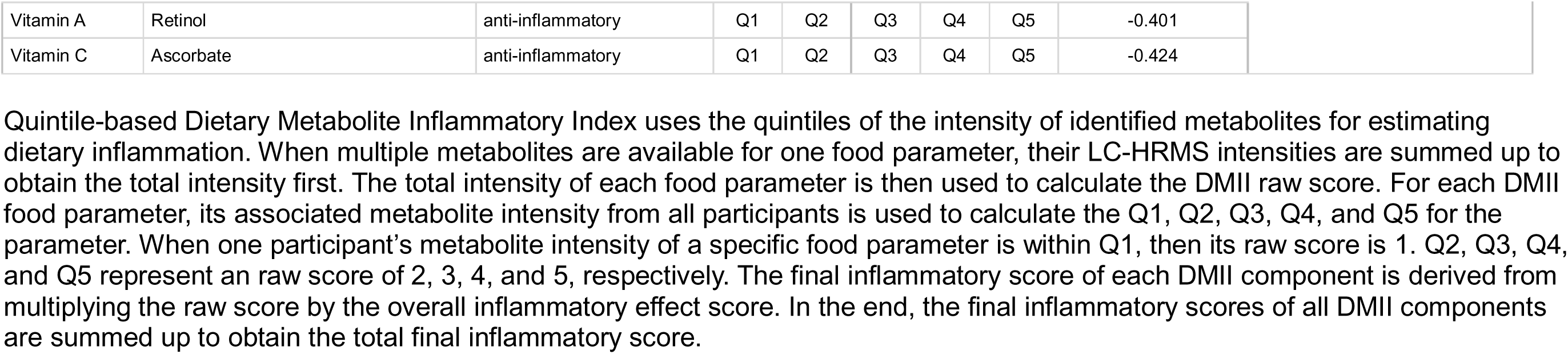
Quintile-based Dietary Metabolite Inflammatory Index.

| Food parameter | Metabolite parameter | Inflammatory effect | Raw score |  |  |  |  | Overall inflammatory effect score | Final inflammatory score |
| --- | --- | --- | --- | --- | --- | --- | --- | --- | --- |
|  |  |  | 1 | 2 | 3 | 4 | 5 |  |  |
| Vitamin B6 | Pyridoxal (B6) | anti-inflammatory | Q1 | Q2 | Q3 | Q4 | Q5 | -0.365 | Individual raw score *<br>individual inflammatory effect score |
| Caffeine | Caffeine | anti-inflammatory | Q1 | Q2 | Q3 | Q4 | Q5 | -0.110 |  |
| Carbohydrate | Glucose | pro-inflammatory | Q1 | Q2 | Q3 | Q4 | Q5 | 0.097 |  |
| Cholesterol | Cholesterol | pro-inflammatory | Q1 | Q2 | Q3 | Q4 | Q5 | 0.110 |  |
| Fiber | Indole propionate | anti-inflammatory | Q1 | Q2 | Q3 | Q4 | Q5 | -0.663 |  |
| Green/black tea | Hippuric acid | anti-inflammatory | Q1 | Q2 | Q3 | Q4 | Q5 | -0.536 |  |
| MUFA | Palmitoleic acid<br>FA 18:1 (Oleic acid)<br>FA 20:1 (Gondoic acid) | anti-inflammatory | Q1 | Q2 | Q3 | Q4 | Q5 | -0.009 |  |
| Niacin | Nicotinamide (B3) | anti-inflammatory | Q1 | Q2 | Q3 | Q4 | Q5 | -0.246 |  |
| n-3 fatty acid | FA 20:5 (Eicosapentaenoic acid)<br>DHA (Docosahexaenoic acid) | anti-inflammatory | Q1 | Q2 | Q3 | Q4 | Q5 | -0.436 |  |
| n-6 fatty acid | FA 18:2 (Linoleic acid)<br>FA 20:2 (Eicosadienoic acid)<br>FA 20:3 (Homolinoleic acid)<br>FA 20:4 (Arachidonic acid)<br>FA 22:4 (Docosatetraenoic acid) | anti-inflammatory | Q1 | Q2 | Q3 | Q4 | Q5 | -0.159 |  |
| Protein | Delta-valerobetaine | pro-inflammatory | Q1 | Q2 | Q3 | Q4 | Q5 | 0.021 |  |
| PUFA | Gamma-Linolenic acid<br>Alpha-Linolenic acid<br>Docosapentaenoic acid (22n-3)<br>Docosapentaenoic acid (22n-6) | anti-inflammatory | Q1 | Q2 | Q3 | Q4 | Q5 | -0.337 |  |
| Saturated fat | FA 14:0 (Myristate)<br>FA 16:0 (Palmitate)<br>FA 17:0 (Heptadecanoic)<br>FA 18:0 (Stearic acid)<br>FA 20:0 (Arachidic acid) | pro-inflammatory | Q1 | Q2 | Q3 | Q4 | Q5 | 0.373 |  |

|  |  |  |  |  |  |  |  |  |
| --- | --- | --- | --- | --- | --- | --- | --- | --- |
| Vitamin A | Retinol | anti-inflammatory | Q1 | Q2 | Q3 | Q4 | Q5 | -0.401 |
| Vitamin C | Ascorbate | anti-inflammatory | Q1 | Q2 | Q3 | Q4 | Q5 | -0.424 |

Quintile-based Dietary Metabolite Inflammatory Index uses the quintiles of the intensity of identified metabolites for estimating dietary inflammation. When multiple metabolites are available for one food parameter, their LC-HRMS intensities are summed up to obtain the total intensity first. The total intensity of each food parameter is then used to calculate the DMII raw score. For each DMII food parameter, its associated metabolite intensity from all participants is used to calculate the Q1, Q2, Q3, Q4, and Q5 for the parameter. When one participant's metabolite intensity of a specific food parameter is within Q1, then its raw score is 1. Q2, Q3, Q4, and Q5 represent an raw score of 2, 3, 4, and 5, respectively. The final inflammatory score of each DMII component is derived from multiplying the raw score by the overall inflammatory effect score. In the end, the final inflammatory scores of all DMII components are summed up to obtain the total final inflammatory score.

### DMII Validation

In the CHDWB study, body mass index, race, HDL cholesterol, triglycerides, fasting glucose, waist-to-hip ratio, hs-CRP, IL-6, and TNF-α differed significantly across the MHNO, MHO, and MUO groups (**Table 4**). In contrast, the DII did not differ significantly across these groups. We first evaluated whether the concentration-based DMII was associated with the FFQ-derived DII in the CHDWB study. The concentration-based (**Figure 4A**), median-based (**Figure 4B**), and quintile-based DMII (**Figure 4C)** were positively correlated with the DII (Figure 4A: Spearman r (r_s_) = 0.28, p = 3.289e-05, Figure 4B: r_s_ = 0.28, p = 0.003, Figure 4C: r_s_ = 0.20, p = 0.004). Concentration-based DMII was strongly correlated with the median-based DMII (r_s_ = 0.62, p = 3.787e-24; **Figure 5A**) and the quintile-based DMII (r_s_ = 0.69, p = 5.002e-31; **Figure 5B**).

**Figure 4.**
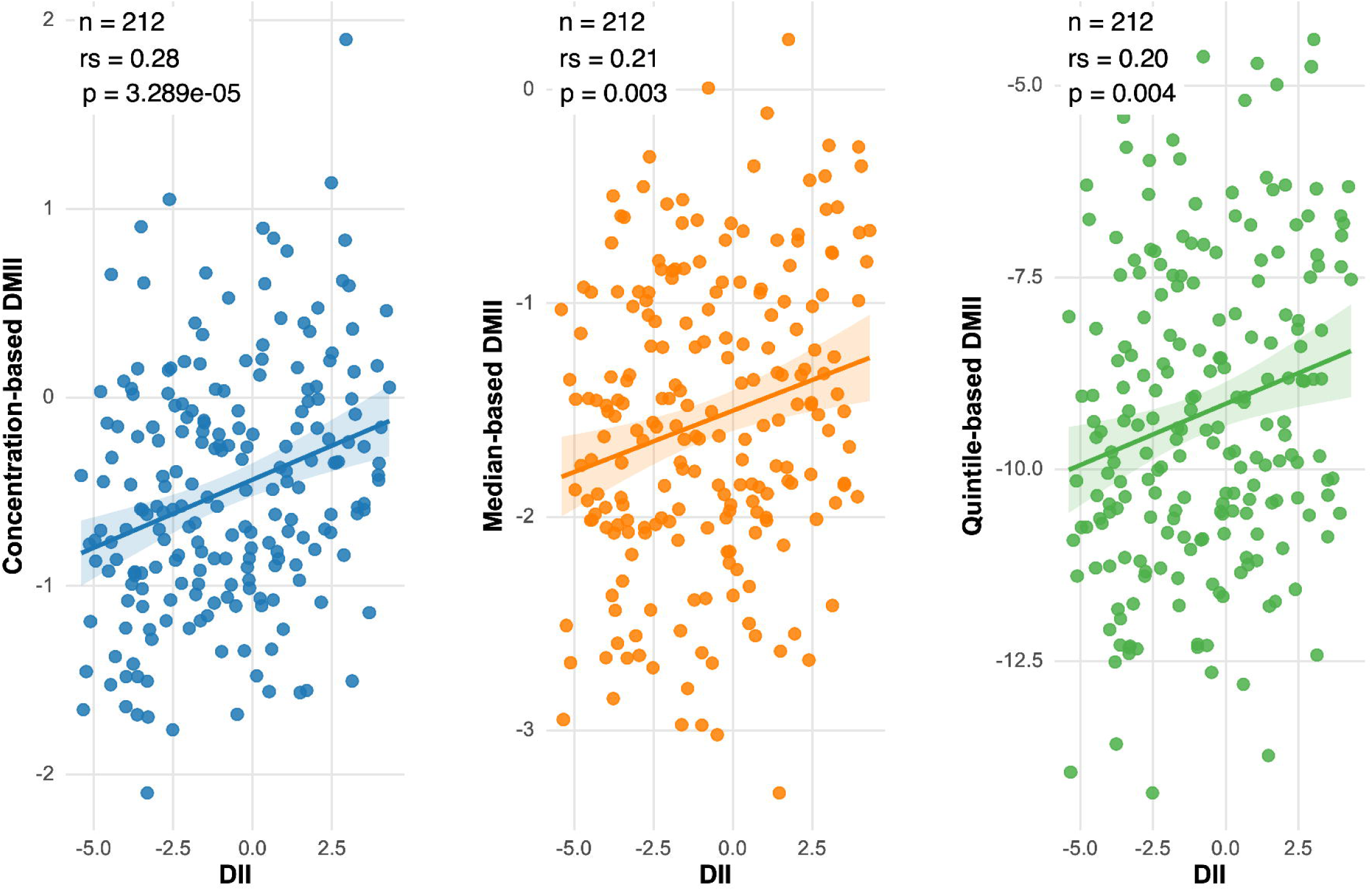
Spearman correlation between the Dietary Inflammatory Index and the Dietary Metabolite Inflammatory Index. Spearman correlation was used for between Dietary Inflammatory Index and different versions of DMII. Only complete observations were used. The Dietary Inflammatory Index was calculated using the *dietaryindex* R package with the Block Food Frequency Questionnaire data. The DMII scores were calculated using the procedure described in Figure 1 and the method section. r_s_ represents Spearman correlation.

**Figure 5.**
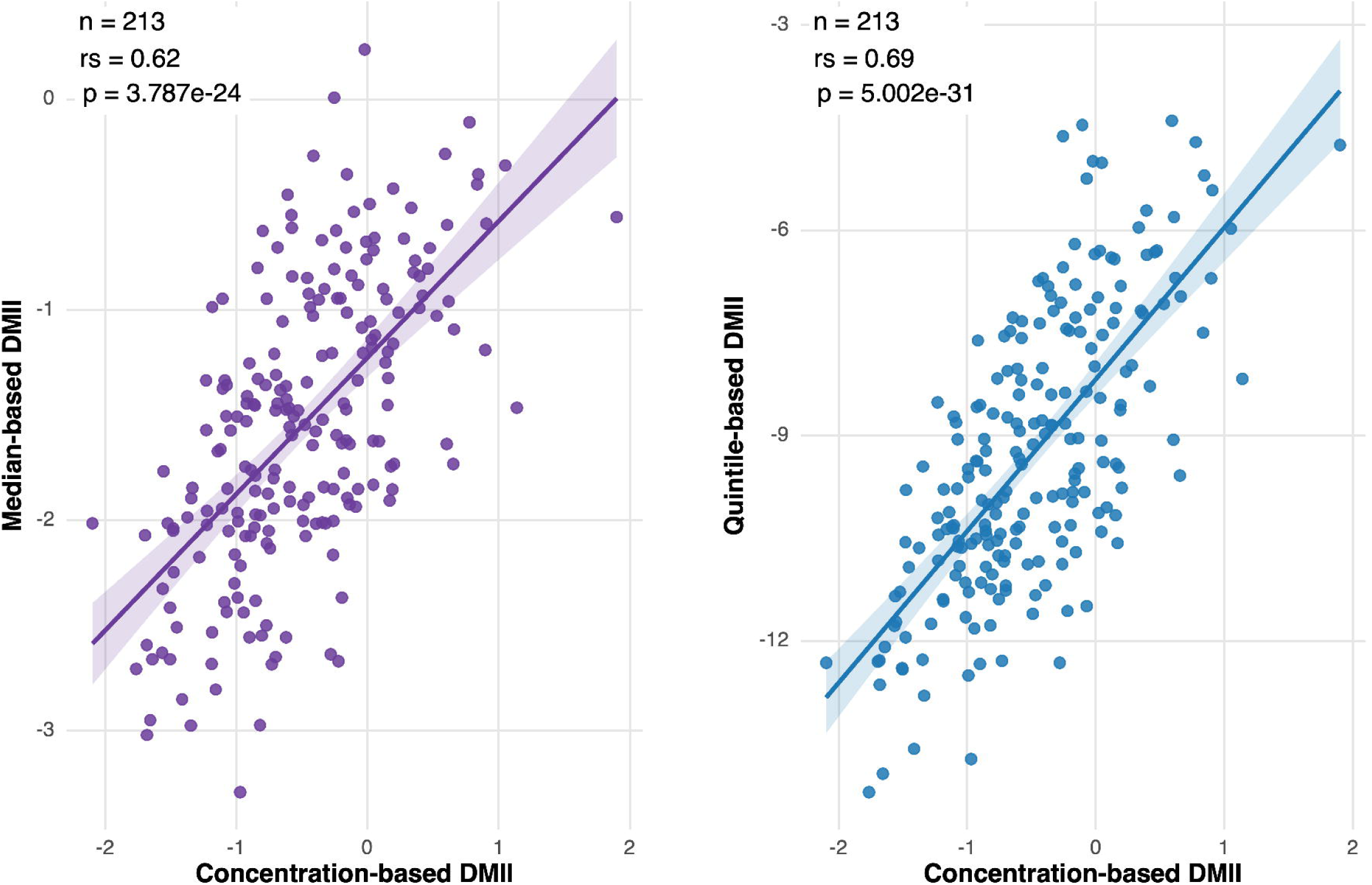
Spearman correlations between different versions of the Dietary Metabolite Inflammatory Index. Spearman correlation was used for the concentration-based DMII, the median-based DMII, and the quintile-based DMII. Only complete observations were used. The DIII scores were calculated using the procedures described in Figure 1 and the method section. r_s_ represents Spearman correlation.

**Table 4.** Demographic Characteristics of the CHDWB Study.

| Group | MHNO<br>(n = 83) | MHO<br>(n = 68) | MUO<br>(n = 61) | P value |
| --- | --- | --- | --- | --- |
| Age | 53.7 ± 4.9 | 52.3 ± 7.1 | 55.1 ± 6.3 | 0.058 |
| Body Mass Index (kg/m<sup>2</sup>) | 23.8 ± 3.1 | 34.8 ± 5.1 | 36.7 ± 5.4 | **1E-39** |
| Sex | 20 (24%) | 16 (24%) | 17 (28%) | 0.826 |
| <u>Race</u> |  |  |  |  |
| Asian | 9 (11%) | 0 (0%) | 1 (2%) | **2E-4** |
| Black | 18 (22%) | 31 (46%) | 13 (22%) | - |
| White | 55 (67%) | 36 (54%) | 44 (76%) | - |
| HDL-Cholesterol (mg/dL) | 73.1 ± 18.1 | 62.2 ± 15.6 | 49.5 ± 13.8 | **3E-14** |
| Triglyceride (mg/dL) | 71 [61–96] | 90.5 [67.2–109.2] | 149 [106–191] | **9E-17** |
| Glucose (mg/dL) | 85 [80–89] | 87 [80.8–92] | 99 [92–109] | **5E-15** |
| Waist-to-Hip Ratio | 0.8 ± 0.1 | 0.8 ± 0.1 | 0.9 ± 0.1 | **4E-11** |
| High sensitivity C-Reactive Protein (mg/L) | 0 [0–0.1] | 0.4 [0.2–0.6] | 0.4 [0.2–0.7] | **2E-19** |
| IL-6 (pg/mL) | 0.5 [0.1–1.1] | 1.4 [0.8–2] | 1.4 [0.7–3] | **3E-5** |
| TNF-α (ng/mL) | 2.7 [1.5–4.3] | 3.9 [2.4–4.5] | 4 [2.6–5.7] | **0.018** |
| Dietary Inflammatory Index Total Score | -0.7 ± 2.5 | -0.6 ± 2.6 | -1.2 ± 2.6 | 0.417 |

MHNO metabolically healthy nonobese, MHO metabolically healthy obese, and MUO metabolically unhealthy obese. Normally distributed continuous variables were compared using Welch's one-way ANOVA; non-normally distributed variables were compared using the Kruskal-Wallis test. Omnibus P values are reported; pairwise post-hoc comparisons were not performed. Categorical variables were compared using Chi-squared tests, or Fisher's exact tests when any expected cell count was less than 5 (Cochran criterion). Statistical significance was defined as $p \leq 0.05$ .

We next tested the DII and the three DMII versions against inflammatory biomarkers in the CHDWB study. The DII was not significantly correlated with hs-CRP, IL-6, or TNF-α (**Table 5**). In contrast, the concentration-based, median-based, and quintile-based DMII were all positively correlated with hs-CRP and IL-6 but not TNF-α (**Table 5**). Metabolites used in the DMII calculations in the CHDWB study are available in **Supplementary Table 2**.

**Table 5.** Spearman Correlations between Dietary Inflammatory Index, Dietary Metabolite Inflammatory Index, and Inflammatory Biomarkers in the CHDWB study.

| Variable 1 | Variable 2 | n | $r_s$ | p-value |
| --- | --- | --- | --- | --- |
| DII | hs-CRP | 212 | 0.06 | 0.384 |
| DII | IL-6 | 153 | 0.031 | 0.701 |
| DII | TNF-α | 161 | -0.119 | 0.133 |
| Concentration-based DMII | hs-CRP | 213 | 0.153 | **0.026** |
| Concentration-based DMII | IL-6 | 153 | 0.183 | **0.024** |
| Concentration-based DMII | TNF-α | 162 | 0.118 | 0.134 |
| Median-based DMII | hs-CRP | 213 | 0.201 | **0.003** |
| Median-based DMII | IL-6 | 153 | 0.231 | **0.004** |
| Median-based DMII | TNF-α | 162 | 0.044 | 0.578 |
| Quintile-based DMII | hs-CRP | 213 | 0.25 | **2.25E-04** |
| Quintile-based DMII | IL-6 | 153 | 0.301 | **1.58E-04** |
| Quintile-based DMII | TNF-α | 162 | 0.016 | 0.843 |

DII Dietary Inflammatory Index, DMII Dietary Metabolite Inflammatory Index, hs-CRP high sensitivity C-reactive protein, $r_s$ Spearman correlation coefficient. Complete observations for each biomarker were used for statistical analysis. Statistical significance was defined as $p < 0.05$ .

We performed an additional validation analysis in the ATLAA study using CRP and IL-6. The DII was not significantly correlated with either biomarker. In contrast, the concentration-based DMII was positively correlated with CRP (r_s_ = 0.144, p = 5.32e-04) and IL-6 (r_s_ = 0.085, p = 0.038) (**Table 6**). Median-based DMII did not have significant correlations with CRP and IL-6. The quintile-based DMII shared a trend toward positive correlation with CRP (r_s_ = 0.081, p = 0.051) (**Table 6**). Metabolites used in the DMII calculations in the ATLAA study are available in **Supplementary Table 3**.

**Table 6.** Spearman Correlations between Dietary Inflammatory Index, Dietary Metabolite Inflammatory Index, and Inflammatory Biomarkers in the ATLAA study.

| Variable 1 | Variable 2 | n | $r_s$ | p-value |
| --- | --- | --- | --- | --- |
| DII | CRP | 450 | 0.019 | 0.685 |
| DII | IL-6 | 474 | 0.055 | 0.231 |
| Concentration-based DMII | CRP | 574 | 0.144 | **5.32E-04** |
| Concentration-based DMII | IL-6 | 598 | 0.085 | **0.038** |
| Median-based DMII | CRP | 574 | 0.031 | 0.458 |
| Median-based DMII | IL-6 | 598 | 0.032 | 0.435 |
| Quintile-based DMII | CRP | 574 | 0.081 | 0.051 |
| Quintile-based DMII | IL-6 | 598 | 0.051 | 0.217 |

DII Dietary Inflammatory Index, DMII Dietary Metabolite Inflammatory Index, CRP C-reactive protein, $r_s$ Spearman correlation coefficient. Complete observations for each biomarker were used for statistical analysis. Statistical significance was defined as $p < 0.05$ .

### DMII application

We then applied the median-based DMII to four Alzheimer’s disease datasets because the ST000046 study only has 15 AD patients and 15 healthy controls and metabolites were not quantified. In each dataset, the odds ratio (OR) represents the association with Alzheimer’s disease per one study-specific standard deviation higher median-based DMII.

The ORs were 1.26 (95% CI: 0.98–1.61) in WHICAP, 1.34 (95% CI: 1.11–1.62) in EFIGA, 0.88 (95% CI: 0.43–1.83) in ST000046 plasma, and 2.63 (95% CI: 1.07–6.46) in ST000046 cerebrospinal fluid (**Figure 6**). The pooled summary OR was 1.31 (95% CI: 1.13–1.52) for the median-based DMII associated with Alzheimer’s disease.

**Figure 6.**
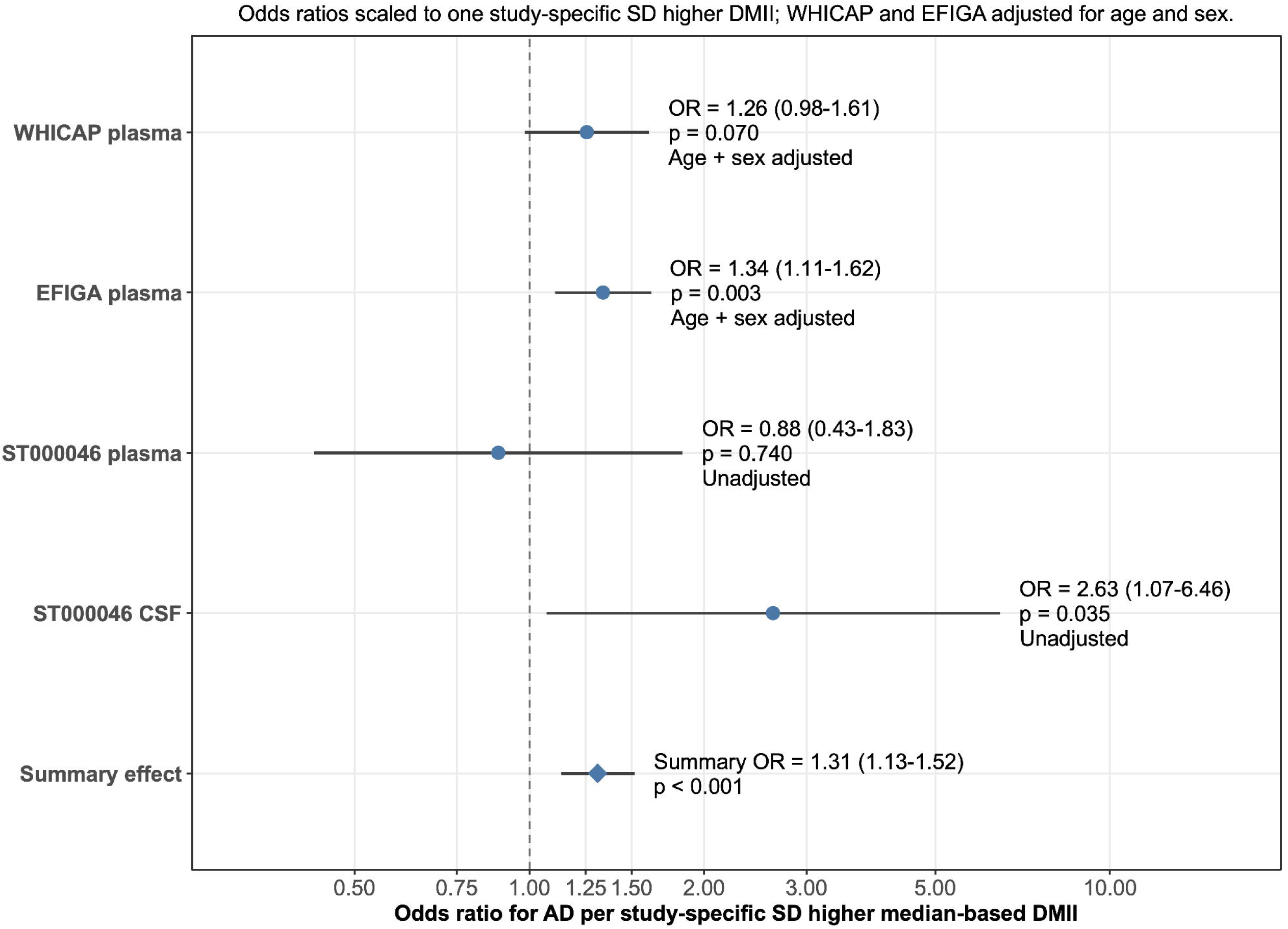
Application of DMII for the meta-analysis of Alzheimer’s disease studies. Odds ratios scaled to one study-specific standard deviation higher median-based DMII; The WHICAP and EFIGA studies’ results were adjusted for age and sex, while ST000046 were not adjusted, as age and sex data were not available.

## Discussion

We developed, tested, and applied three versions of Dietary Metabolite Inflammatory Index (DMII) to assess diet-related chronic inflammation using LC-HRMS untargeted metabolomics data. These included concentration-based, median-based, and quintile-based DMII scores. All three versions of the DMII score showed significant but weak correlations with the DII. These weak correlations may reflect interindividual differences in metabolism, including genetic variation and differences in nutrient absorption, metabolism, and disposition, as well as different numbers of scoring components. The three DMII versions were also significantly correlated with each other, suggesting that they captured related diet-related inflammation as measured by metabolomics.

In the validation analyses, the DMII scores were significantly correlated with hs-CRP, CRP, and IL-6, whereas the DII was not. This difference may reflect the complementary strengths of metabolomics-based dietary assessment. LC-HRMS untargeted metabolomics measures dietary metabolites directly in human biospecimens and can capture variations in genetics and nutrient absorption, metabolism, and disposition. In contrast, the Block FFQ depends on self-reported food consumption frequency and quantity and is therefore measured nominal dietary intake affected by recall and reporting bias. Thus, the DMII may provide a biologically relevant measure of diet-related inflammation that complements a questionnaire-based dietary index.

The strength of the associations between DMII scores and inflammatory biomarkers differed between cohorts. These correlations were stronger in the CHDWB study than in the ATLAA study, even though both cohorts included participants with obesity who were at high risk of chronic inflammation. One possible explanation is that the ATLAA study included pregnant women. Pregnancy can alter circulating levels of inflammatory biomarkers, which may have weakened or modified the observed associations between DMII scores and inflammation (50,51).

We then applied the DMII to four publicly available Alzheimer’s disease metabolomics datasets as a proof-of-concept application. These datasets did not have dietary assessment data but had LC-HRMS untargeted metabolomics data. The DMII results were consistent with previous studies linking dietary inflammation to Alzheimer’s disease risk (15,52,53). Interestingly, the ST000046 cerebrospinal fluid dataset showed a stronger association with Alzheimer’s disease than the paired plasma dataset, even though the two datasets included the same participants. The odds ratio was 2.63 (95% CI: 1.07–6.46) in cerebrospinal fluid and 0.88 (95% CI: 0.43–1.83) in plasma, which is perhaps due to the small sample size.

Several dietary indexes are available to assess dietary patterns, and precision nutrition is an emerging field. However, no standardized tool is available to assess diet-related inflammation using metabolomics data. To our knowledge, the DMII is the first standardized approach for calculating diet-related inflammation using either quantified or unquantified metabolites from untargeted metabolomics datasets. This approach allows researchers to calculate diet-related inflammatory potential in clinical and epidemiological studies without requiring traditional dietary assessment data. In principle, this therefore provides a new foundation for predicting diet-related inflammation in clinical nutrition practices.

Three versions of the DMII are available so that researchers can select the version that best fits their study design and data type. The concentration-based DMII requires metabolite quantification. Its main advantage is that it can support comparisons across studies because it uses standardized metabolite concentrations. It may also support clinical nutrition applications because the concentration-based DMII can be calculated from a single blood sample. The DMII also provides a framework for studying mechanisms of diet-related chronic inflammation. When the DMII differs between disease and control groups, researchers can perform metabolome-wide association analyses to identify metabolic alterations. These analyses can examine which DMII component metabolites differ by disease status, how downstream metabolites are altered, and which metabolic pathways overlap between DMII-associated changes and disease-associated changes.

The median-based and quintile-based DMII were developed for untargeted metabolomics studies without absolute metabolite quantification. The median-based DMII is more flexible and may be suitable for cross-sectional studies with small sample sizes, especially when each study group includes fewer than 40–50 participants. The quintile-based DMII also supports unquantified metabolomics data, but it requires a larger sample size. In general, each study group should include at least 50 participants to support reliable quintile classification and statistical analysis. Quintile-based DMII may work better than median-based DMII when the sample size is sufficient, since it uses more granular scoring criteria for DMII.

Several limitations should be considered. First, the current DMII does not include metabolites related to several DII food and nutrient parameters, as they were not identified and quantified by our LC-HRMS platform. These include flavonoids, such as flavan-3-ols, flavones, flavonols, flavanones, anthocyanidins, and isoflavones; minerals, such as iron, magnesium, selenium, and zinc; antioxidant-rich foods and spices, such as thyme, oregano, rosemary, eugenol, garlic, ginger, onion, saffron, turmeric, and pepper; vitamins, such as vitamin B12, vitamin D, and vitamin E; and other parameters, including beta-carotene, alcohol, energy, total fat, and trans fat. Although identified but not quantified candidate metabolites were available for some of these parameters in median- and quintile-based DMII, including them weakened the correlations (data not shown) with the inflammatory biomarkers. This may reflect the metabolism and disposition of the metabolites in blood. Therefore, these metabolites were not included in the final DMII. The second limitation is that some DMII metabolites are not specific to a single food source. For example, caffeine can reflect dietary intakes of coffee, tea, or energy drink. Hippuric acid can reflect dietary intakes of fruits, vegetables, coffee, cocoa, or tea (54). However, this limitation is not unique to the DMII. Dietary indexes may include parameters that are not mutually exclusive (2). Therefore, the limited specificity of individual metabolites may not prevent the DMII from capturing broad diet-related chronic inflammation. Third, the construction of the DMII relies on the sample processing and spectra acquisition of LC-HRMS untargeted metabolomics data. If the metabolomics data missed certain metabolites in the DMII, DMII may underperform. Fourth, the DMII correlation analyses do not account for interindividual genetic differences and environmental toxicant exposures, even though these factors have known associations with chronic inflammation (55–57).

To facilitate DMII calculation, we developed the MetaboIndex R package (https://github.com/jamesjiadazhan/MetaboIndex) that is available on GitHub. This package supports metabolite reference standardization, metabolite identification using reference libraries or the MSMICA algorithm, and DMII score calculation using quantified or unquantified metabolites. The DMII may be especially useful when dietary data are unavailable, when dietary sample size is small to moderate, when a precise measure of diet-related inflammation is needed for an individual, or when researchers want to analyze archived untargeted metabolomics datasets for nutrition-related questions.

## Author contributions

The authors’ responsibilities were as follows – JZ: designed the research; JZ, CY: calculated the Dietary Inflammatory Index; JZ: designed and calculated the Dietary Metabolite Inflammatory Index; YT, DL, AD: provided access to the metabolomics and inflammatory datasets of the ATLAA study; GM, DPJ: provided access to the metabolomics and inflammatory datasets of the CHDWB study; JZ: conducted the data analysis; JZ: completed the initial draft of the manuscript; JZ, CY, YT, RS, JA, DL, AD, GM, YG, DPJ: revised the manuscript; JZ and DPJ had primary responsibility for the final content; and all authors: read and approved the final manuscript.

## Funding

The corresponding author received funding from the following sources: NIH/NIA R21 AG080247 and R01 AG085279.

## Data Availability

Raw metabolomics, inflammatory biomarker, and dietary data of the CHDWB study will be uploaded to the metabolomics workbench.

## Conflict of interest

J.Z. and D.P.J. are listed as inventors on a U.S. provisional patent application filed by Emory University related to the technology described in this manuscript. The patent application is managed by Emory University Office of Technology Transfer. If the technology is licensed or commercialized, the inventors may receive revenue according to Emory University policies. The authors declare no other competing interests.

## Notes

https://github.com/jamesjiadazhan/MetaboIndex

